# Sexual selection rewires reproductive protein networks

**DOI:** 10.1101/422410

**Authors:** Timothy L. Karr, Helen Southern, Matthew Rosenow, Toni I. Gossmann, Rhonda R. Snook

**Author notes:** Corresponding authors: &. Current address: Arizona State University, School of Life Sciences, Tempe, AZ 85287.

## Abstract

Polyandry drives postcopulatory sexual selection (PCSS), resulting in rapid evolution of male ejaculate traits. Critical to male and female fitness, the ejaculate is known to contain rapidly evolving seminal fluid proteins (SFPs) produced by specialized male secretory accessory glands. The evidence that rapid evolution of some SFPs is driven by PCSS, however, is indirect, based on either plastic responses to changes in the sexual selection environment or correlative macroevolutionary patterns. Moreover, such studies focus on SFPs that represent but a small component of the accessory gland proteome. Neither how SFPs function with other reproductive proteins, nor how PCSS influences the underlying secretory tissue adaptations and content of the accessory gland, has been addressed at the level of the proteome. Here we directly test the hypothesis that PCSS results in rapid evolution of the entire male accessory gland proteome and protein networks by taking a system-level approach, combining divergent experimental evolution of PCSS in *Drosophila pseudoobscura (*Dpse), high resolution mass spectrometry (MS) and proteomic discovery, bioinformatics and population genetic analyses. We demonstrate that PCSS influences the abundance of over 200 accessory gland proteins, including SFPs. A small but significant number of these proteins display molecular signatures of positive selection. Divergent PCSS also results in fundamental and remarkably compartmentalized evolution of accessory gland protein networks in which males subjected to strong PCSS invest in protein networks that serve to increase protein production whereas males subjected to relaxed PCSS alters protein networks involved in protein surveillance and quality. These results directly demonstrate that PCSS is a key evolutionary driver that shapes not only individual reproductive proteins, but rewires entire reproductive protein networks.

**The abbreviations used are:**

BLAST: Basic Local Alignment Search Tool
Dpse: *Drosophila pseudoobscura*
PCSS: postcopulatory sexual selection
SFPs: seminal fluid proteins
Dmel: *D. melanogaster*
SDS: sodium dodecylsulfate
SDS-PAGE: sodium dodecylsulfate polyacrylamide gel electrophoresis
MS: mass spectrometry
LC-MS/MS: liquid chromatography-MS/MS
AcgP: accessory gland proteome
FDRs: False Discovery Rates
AcgS: accessory gland secretome
exoP: exoproteome
LFQ: label-free quantitation
P: polyandry
M: monandry
GO: gene ontology
CC: cellular component
MF: molecular function
BP: biological process
STRING: Search Tool for the Retrieval of Interacting Genes/Proteins
DIOPT: DRSC Integrative Ortholog Prediction Tools
ER: endoplasmic reticulum

---

Polyandry, in which females mate with different males across a reproductive bout, generates PCSS in which ejaculates compete for fertilization of a limited supply of ova and females may choose whose sperm will fertilize those limited ova (1). Polyandy also engenders sexual conflict, in which male and female reproductive interests differ, as a consequence of the disproportionate costs and benefits of mating between the sexes (2). PCSS operates in internally fertilizing organisms between the female reproductive tract and the male ejaculate, which is composed of both sperm and non-sperm components, including SFPs (3). SFPs were first identified by their canonical signal peptide sequence that direct proteins to the secretory pathway (4). Cross-species comparative work has found that general classes of SFPs are conserved (e.g., proteases and protease inhibitors, lectins and prohormones) suggesting that their mechanisms of action are also conserved. However, individual SFPs can rapidly evolve with signals of accelerated rates of adaptive molecular evolution found in studies of coding sequence and male-biased gene expression observed across different animal taxa (e.g., mammals (5, 6); birds (7); Drosophila (8–10)).

Rapid evolution of SFPs is attributed to PCSS because this cocktail of proteins, transferred to females, has dramatic and sometimes antagonistic effects on male and female fitness, including increased female fecundity, reduced female receptivity, decreased female life span, and remodelling of female reproductive tract morphology (4, 11–13). Moreover, males can tailor ejaculate composition, altering the quantity of specific SFPs, via plastic changes in response to variation in the PCSS environment (5, 14) although a mechanism by which males accomplish this is unknown. However, while many studies have examined the evolution and fitness consequences of SFPs, most of them examine the impact of one or just a few of these proteins. This focus is problematic because SFPs do not work in isolation, but rather as partners in protein-protein and protein-nucleic acid networks, and not just solely with other SFPs (15, 16). Therefore, changes in either protein levels or protein function may have significant up-and/or down-stream consequences on organismal physiology and fitness. Indirect studies of the role of PCSS on reproductive protein evolution have not considered other proteins also produced by male accessory gland tissue that may interact with SFPs and influence fitness.

Most of our current knowledge of SFPs comes from traditional protein detection methods and comparisons aimed at one species, *D. melanogaster* (9, 17–19). However, the advent of sophisticated transcriptomic, proteomic and bioinformatics approaches have identified new SFPs and associated reproductive proteins not only in Dmel but also other taxa (20–26). Furthermore, high throughput mass spectrometry based techniques now routinely produce cellular and tissue proteomes consisting of thousands of protein IDs not only useful for protein identification but also the analysis and identification of protein interaction networks.

Initial application using these techniques to the analysis of the sperm proteome (24, 25) found heterogeneity in the evolutionary rates of genes compared to SFPs which lead to the hypothesis that adaptive compartmentalization can occur both within and between proteomes involved in reproduction. This hypothesis suggests that interacting reproductive proteins would comprise a set of “core” proteins with essential cellular functions (e.g., sperm motility) that are predicted to be under strong purifying selection and another set of proteins under adaptive selection generated from PCSS, e.g., sperm-egg interactions (24). Other than sperm, direct empirical confirmation of adaptive compartmentalization, and a possible role of PCSS in this process, has yet to be undertaken. Likewise, application of high throughput mass spectrometry based techniques have allowed characterization and dynamics of protein interaction networks in a variety of cellular contexts including disease-induced perturbation and or over evolutionary time frames (see (27, 28) for recent reviews). However, the role of PCSS in shaping the architecture of protein interaction networks is unknown.

Here we address the overall qualitative and quantitative impact of PCSS on the male accessory gland proteome and directly test the hypothesis that PCSS effects SFP production and their reproductive protein networks. We also test for signatures of positive selection and examine evidence for adaptive compartmentalization. We perform these tests using replicated experimentally evolved lines of *D. pseudoobscura* (Dpse) in which PCSS was manipulated over many generations by either allowing polyandry (“P” lines; 1 female with 6 males) or enforcing monogamy (“M” lines; 1 female with 1 male) (29). Previous studies using these lines have found that divergent sexual selection impacts the evolution of reproductive traits relevant to PCSS and SFP production, including P males altering female investment in both early and total progeny production (30) and P males having larger accessory glands that resulted in greater reproductive success (31). Sex-biased gene expression has also evolved in these lines (32, 33), and, taken togther these results strongly motivate using these lines to directly test the role of PCSS in shaping male reproductive proteomes at the microevolutionary scale.

## EXPERIMENTAL PROCEDURES

### Experimental lines

The establishment and maintenance of the selection lines were previously described in detail (29) Briefly, an ancestral wild-caught population of Dpse from Tucson Arizona USA, a naturally polyandrous species (wild caught females have been shown to be frequently inseminated by at least two males at any given time; (34)), was used to establish the selection lines. From this population, four replicate lines (replicates 1-4) of two different sexual selection treatments were established. To modify the opportunity for sexual selection, adult sex ratio in vials was manipulated by either confining one female with a single male and enforcing monogamy (“monogamy” treatment, M) or one female with six males promoting polyandry (“polyandry” treatment, P (NB: this treatment has also been referred to as E in other publications). Effective population sizes are equalized between the treatments as described previously (35). At each generation, offspring are collected and pooled together for each replicate line, and a random sample from this pool is used to constitute the next generation in the appropriate sex ratios, thus proportionally reflecting the differential offspring production across families. In total, eight selection lines (M1, M2, M3, M4 and P1, P2, P3, P4) are maintained, in standard vials (2.5 × 80 mm), with a generation time of 28 days. All populations are kept at 22°C on a 12L:12D cycle, with standard food media and added live yeast.

### Experimental flies

Flies from replicates 1-4 of each of the selection lines were collected from generations 157, 156, 155 and 153 respectively. We standardized for maternal and larval environments (29), but in brief, parental flies were collected and housed en-mass in food bottles, then groups of about 30 were transferred on egg laying plates for 24 hours, removed and replaced with a fresh egg plate. This second plate was removed after 24 hours, then 48 hours later, first instar larvae were collected in groups of 100 and housed in standard molasses/agar food vials at 22°C. Males from these vials were collected on the day of eclosion and housed in vials of 10 individuals, until they were sexually mature (36), and then dissected when they were 5 or 6 days old.

### Accessory gland tissue preparation

For dissections, males were anaesthetized with ether, placed in a drop of phosphate buffered saline and the reproductive tract removed. Intact accessory glands were clipped from the rest of the reproductive tract (Supplemental Figure 1A, B) with fine dissection needles, moved to another drop of phosphate buffered saline for rinsing, and then transferred to a microcentrifuge tube. 30 accessory glands per replicate were used for subsequent LC-MS/MS as described below. Samples from each replicate were solubilized at 4°C by addition of 30 μl of RIPA buffer (Sigma), and a HALT protease inhibitor mixture containing phenylmethylsulfonyl fluoride (Thermo Fisher Scientific). Once all accessory glands had been dissected, samples were freeze/thawed three times on dry ice (∼5 mins) and then thawed at 37°C for 30 seconds. After the freeze/thaw cycles, samples were vortexed for 30 seconds, centrifuged at 20,000 rpm for 5 minutes at 4°C and then stored at −80°C until processing.

### Sodium dodecylsulfate polyacrylamide gel electrophoresis (SDS-PAGE) and in-gel digestion of proteins

Protein concentration was determined using a Bradford assay and samples were solubilized in SDS sample buffer containing 10 mM dithiothreitol and proteins separated on 4-12% SDS-PAGE gels per manufacturer instructions (Invitrogen). Protein bands were visualized (Supplemental Figure 1C) using Brilliant Blue G Colloidal Concentrate (Sigma). Each gel lane was manually cut into approximately equivalent sized pieces and destained using 200 mM ammonium bicarbonate and 40% acetonitrile. Gel pieces were then reduced in 200 μl of a 50 mM ammonium bicarbonate buffer containing 10 mM dithiothreitol, followed by alkylation in a similar volume of a 50 mM ammonium bicarbonate containing 55 mM iodoacetamide. Gel pieces were then centrifuged at 13 Kg for 10 seconds and dried using a vacuum concentrator until all samples were dry (∼30 min). The dried pieces were then hydrated in a solution containing 20 μl of trypsin (New England BioLabs) and 50 μl of acetonitrile and incubated overnight at 37°C. Peptides were extracted the following day using a standard method with a solution of 100% acetonitrile and 5% formic acid and dried down overnight in a vacuum concentrator at 30°C. Resulting peptides were resuspended in 7.5 μl of 0.1% (v/v) formic acid, 3% (v/v) acetonitrile, sonicated in a water bath for 5 minutes and centrifuged at 13 × g for 10 seconds, before being transferred to a sample vial and loaded into the autosampler tray of the Dionex Ultimate 3000 μHPLC system. Samples were set to run using the Xcalibur sequence system.

### Liquid chromatography-MS/MS (LC-MS/MS) data collection

All MS data were collected using an LTQ Orbitrap Elite hybrid mass spectrometer (Thermo Fisher Scientific) equipped with an Easy-Spray (Thermo Fisher Scientific) ion source. Peptides were separated using an Ultimate 3000 Nano LC System (Dionex). Peptides were desalted on-line using a capillary trap column (Acclaim Pepmap100, 100 μm, 75 μm × 2 cm, C18, 5 μm; Thermo Fisher Scientific) and then separated using 60 min reverse phase gradient (3-40% acetonitrile/0.1% formic acid) on an Acclaim PepMap100 RSLC C18 analytical column (2 μm, 75 μm id × 10 cm; Thermo Fisher Scientific) with a flow rate of 0.25 μl/min. The mass spectrometer was operated in standard data dependent acquisition mode controlled by Xcalibur 2.2. The instrument was operated with a cycle of one MS (in the Orbitrap) acquired at a resolution of 60,000 at m/z 400, with the top 20 most abundant multiply-charged (2+ and higher) ions in a given chromatographic window further subjected to CID fragmentation in the linear ion trap. An FTMS target value of 1e6 and an ion trap MSn target value of 10000 were used. Dynamic exclusion was enabled with a repeat duration of 30 s with an exclusion list of 500 and exclusion duration of 30 s.

### Experimental Design and Statistical Rationale

#### Database construction of the Dpse accessory gland proteome (AcgP)

The mass spectra data files were searched individually using Sequest HT within the Proteome Discover suite (Thermo Fisher Scientific, San Jose, CA, USA; version 1.4.1.14) using *Drosophila pseudoobscura pseudoobscura* fasta file, (December, 2015 release). Peptide matches were further analyzed and validated within Scaffold Q+ (Proteome Software; version 3.2.0) using X!Tandem. Sequest HT and X!Tandem searches were set with a fragment ion mass tolerance of 0.60 Da and a parent ion tolerance of 10.0 parts per million. The oxidation of methionine (15.99), carboxyamidomethyl of cysteine (57.02), and acetyl modification on peptide N-terminus (42.01) were set as variable modifications. Files from Sequest HT searches within the same gel lane were merged together as Mudpit using Scaffold which calculated False Discovery Rates (FDRs) using a reverse concatenated decoy database (FDR was set at 1.0%). Peptide identifications were accepted if they could be established at greater than 95.0% probability as specified by the PeptideProphet^48^ and protein identifications were accepted if they could be established at greater than 99.0% probability and contained at least 2 identified peptides. Protein probabilities were assigned by the Protein Prophet Algorithm (37). Proteins that contained similar peptides and could not be differentiated based on MS/MS analysis alone were grouped to satisfy the principles of parsimony. The dataset was filtered so that every protein must be identified by at least two unique peptides in any one of the biological replicates. Although a conservative approach, this procedure ensured a robust dataset devoid of potential misidentifications often caused by use of a single peptide for protein identification. To establish a working list of the AcgP, protein IDs from Scaffold were converted to Dpse Fly Base gene numbers (FBgns) using the Uniprot website (Uniprot.org). The resulting Dpse FBgns were then used to query Flybase (Flybase.org) to retrieve orthologous Dmel genes from the OrthoDB orthology tables as implemented in Flybase (38). A complete listing can be found in Supplemental Table 1.

#### Database construction of the Dpse accessory gland secretome (AcgS) and exoproteome (exoP)

As a secretory organ, the accessory gland is expected to contain the cellular machinery necessary for efficient and sustained secretory activity throughout the adult reproductive life cycle. To examine and focus on potential activities related to secretion, we assembled an *in silico* AcgS from the 3281 FBgns of the AcgP as input into Uniprot resulting in 5624 UniProtKB IDs (which includes all predicted protein isoforms). Fasta protein sequence files from each Uniprot entry were downloaded and input into SignalP (http://www.cbs.dtu.dk/services/SignalP/) (39) and Phobius (http://phobius.sbc.su.se) (40), using default settings. The protein IDs were combined, exported into Excel, yielding a final list of 771 UniProt identifiers. The Uniprot IDs were mapped back to 535 unique Dpse FBgns (via Uniprot) which, after submission to OrthoDB (via Flybase) resulted in a final list and 506 Dmel orthologs. Candidate SFPs were identified by first querying the list of 515 AcgS genes in Flybase for Gene Ontology (GO) terms containing “extracellular”. The resulting list of 151 proteins therefore represents the accessory gland exoproteome (exoP) and is considered to contain a representative sampling of a major fraction of SFPs.

#### Network and protein interaction network analyses

The finalized datasets were used for downstream bioinformatic analyses and subsequent visualizations of GO enrichment. The protein coding sequences of the AcgP were downloaded from Uniprot and submitted to Blast2go for annotation and tabulation of the three major GO categories, biological process (BP), molecular function (MF) and cellular component (CC). GO enrichment and network visualization and analysis was performed with Cytoscape v3.4 (41, 42) and ClueGO plugin version 2.2.4 (43, 44). Network parameters used were specific for each dataset as detailed in figure legends and supplemental tables. Protein interaction network analysis was performed using Search Tool for the Retrieval of Interacting Genes/Proteins (STRING), a program that calculates the degree of protein-protein network interconnectivity (45). A combined set of differentially abundant proteins from each M-and P-line (Qspec, see next section below) was uploaded to the STRING website and the analysis run using the “high confidence 0.9” setting and the “experimental” and “databases” selected for “evidence”. Proteins in each M-and P-up groups were distinguished by color-code: red (M>P) and green (P>M).

#### Label-free quantitation and statistical analysis

To test for differential abundance of male reproductive proteins in response to postcopulatory sexual selection statistical signifcance was calculated using a Bayesian approach as implemented by Qspec (46), part of the statistical package Qprot (47). Qspec provides a statistics framework that calculates Bayes factors-essentially likelihood measures of significance in the context of a generalized linear mixed effects model. Local implementation of Qprot (http://sourceforge.net/p/qprot/) provided command line access to QSpec that calculated Z-statistics, log-fold change estimates and a local FDR for each pair-wise comparison. This approach has been shown effective in capturing a broad range of differentially abundant proteins using LFQ methods (46, 48). Raw spectral counts of the M and P line datasets from Scaffold were input into Qspec and the output imported into Excel. Differential protein abundance differences were calculated using a 1% false-discovery rate (FDR) cut-off, and proteins assigned based on log-fold changes relative to the M-line, i.e., positive fold-change, M < P; negative fold-change, M > P.

#### Molecular evolution parameters

We obtained coding sequence information for the genomes of the two close relatives of Dpse, *D. lowei* and *D. affinis*, downloaded from http://popoolation.at/lowei_genome and http://popoolation.at/affinis_genome (49). These two species have the same karyotype as Dpse and show reasonable divergence (median pairwise dS=0.102 versus *D. lowei* and dS=0.26 versus *D. affinis*) thus avoiding substitution rate saturation. To identify orthologs of the identified AcgP Dpse proteins in the two other Drosophila genomes we combined two approaches. First we used gene annotation ignoring isoforms specification (as these are difficult to identify within a proteomic screen). We then used best BLAST hits (50) of the Dpse gene against each of the two other genomes, but excluded gene sets for which annotation was contradictory to the Dpse annotation. Using a pipeline we developed earlier (51), sequences were aligned using MUSCLE (52), uncertain sequences filtered out using ZORRO (53) and input files converted with pal2nal (54). Sequences were then analyzed using PAML v4.9 (55) to obtain dN/dS values for each gene set (one-ratio estimates) as well as to conduct direct tests of positive selection as implemented in the site-specific models models M7a and M8a. For the one-ratio estimates, median differences in dN, dS and dN/dS among groups (AcgP, AcgS, exoP and differentially abundant M and P proteins) were tested using a non-parametric two-tailed test (57). For the site tests of positive selection, we obtained significance by conducting LRTs as described in the PAML manual and corrected P-values for multiple testing using an FDR=0.1 using the method of Benjamini and Hochberg (56). Median differences in dN, dS and dN/dS among groups (AcgP, AcgS, ExoP and differentially abundant M and P proteins) were tested using a non-parametric two-tailed test (57). Enrichment between counts per groups were tested using a 2×2 Fisher’s exact test.

## RESULTS

### The Dpse AcgP

A Dpse AcgP was constructed from peptide-based (“bottom-up”) shotgun MS/MS spectral data obtained from eight independent runs consisting of four replicate samples from each of two Dpse experimental sexual selection populations. For the purposes of assembling a proteome with the broadest coverage, data from all runs were pooled together resulting in a total of 3757 UniProt IDs that mapped to 3281 unique FlyBase gene names. The M-and P-datasets were highly correlated with >90% (3534/3757) overlap (Figure 1A). Likewise, proteins with values in all four replicates for each PCSS treatment represented the majority of identified proteins (M-line 2103/3649; 57.6% and P-line, 2235/3642; 61.4%) A complete listing and tabulation of these results can be found in Supplemental Table S1. The small number of proteins unique to each population (M-unique = 115; P-unique = 108; Figure 1A) most likely represent missed protein assignments due to low quantities (as measured by total spectral counts). Indeed, the average total spectral counts for the unique set of proteins (ave. 4.5; n = 223) was 16-fold lower than the average across the entire dataset (ave. 72.8, n = 3874).

**Figure 1.**
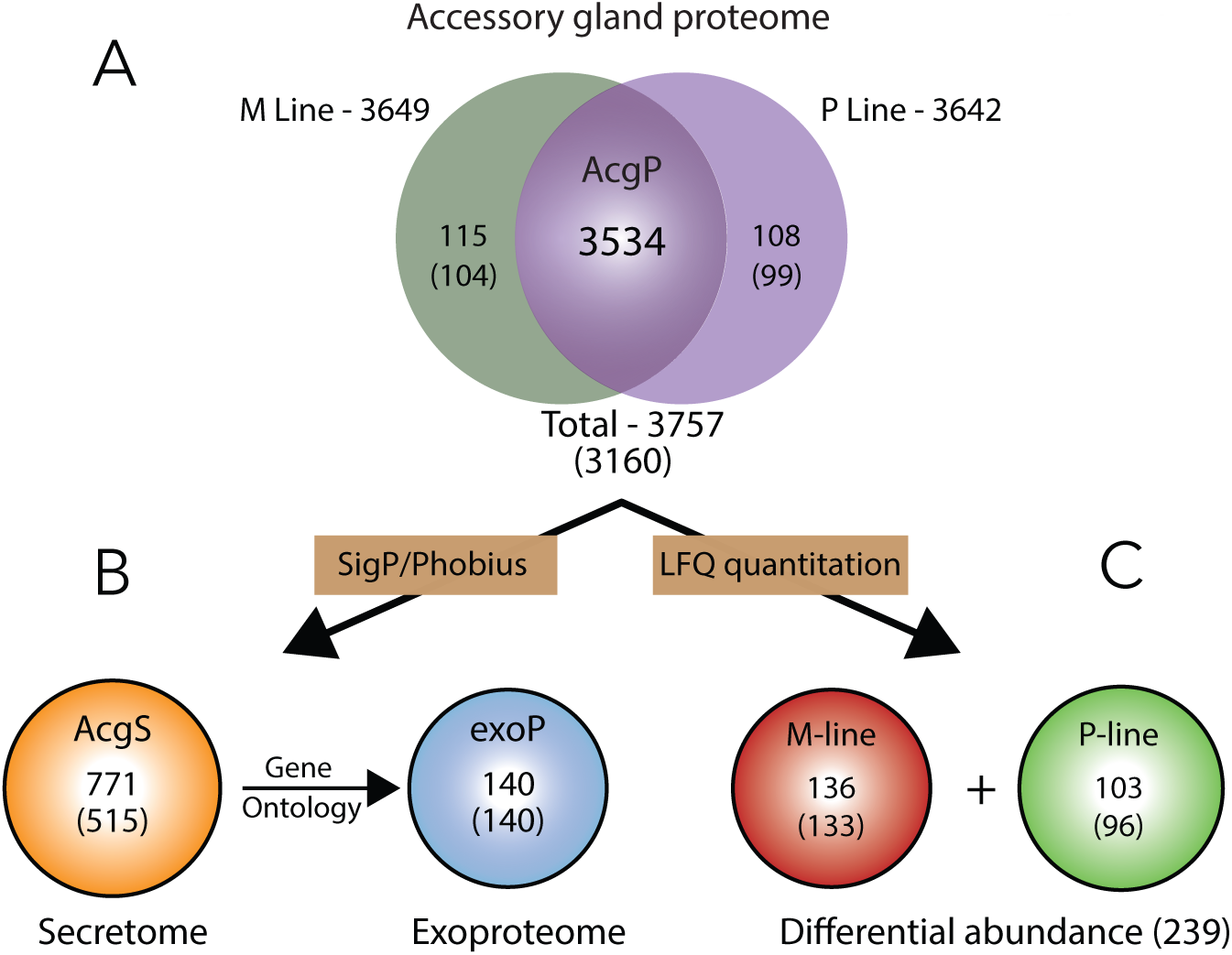
Dpse accessory gland summary statistics. (A) Venn diagram of combined and total Uniprot IDs from the M-and P-experimental lines used to assemble the AcgP. (B) Secretome and exoproteome from the AcgP derived *in silico* (see Methods). (C) Differential protein abundance between the M-and P-experimental lines. Numbers in parentheses indicate Dmel orthologs from OrthoDB database.

### GO and pathway analysis of the AcgP

Pathway and functional analyses began by identifying orthologous Dmel genes, (3159/3281, 96.2%; Supplemental Table 1), providing a useful annotated database to evaluate the overall patterns of the functional elements in the AcgP. As an overview of the major GO groupings, Blast2Go returned “signaling”, “reproduction” and “localization” among BP-and numerous proteins annotated as extracellular involved in the CC-categories, respectively (Supplemental Figure 2). Statistical analysis for GO category enrichment (Figure 2) identified BP terms related to intracellular transport (n = 363, P = 2.80E-45), translation (n = 257, P = 2.47E-35), establishment of protein localization (n= 418, P = 1.22E-44), vesicle-mediated transport (n= 301, P = 8.47E-42), endocytosis (n = 106, P = 7E-10), and secretion (n = 140, P = 2.71E-16). Likewise, the AcgP contains a sgnificant number of proteins in CC categories annotated as “extracellular region” (n = 266, P = 3.2E-05), “endomembrane system” (n = 583, P = 1.04E-57) and “vesicle” (n = 230, P = 8.72E-25). Finally, the overall known biochemical pathways of the AcgP, analyzed using the Kyoto Encyclopedia of Genes and Genomes (KEGG), revealed a similar enrichment of 20 overview terms curated by KEGG that included ribosome biogenesis, protein export, endocytosis and the Wnt signaling pathways (Table 1). We conclude the AcgP contains features expected of a metabolically active cell enriched for protein production and protein secretion. See Supplemental Table 2 for a complete listing of all enriched GO categories.

**Table 1.** Enriched KEGG pathways of the AcgP.

| *GOID* | *GO Term* | *Nr. Genes* | *Pvalue\** |
| --- | --- | --- | --- |
| KEGG:03008 | Ribosome biogenesis | 33 | 3.5E-12 |
| KEGG:03040 | Spliceosome | 82 | 8.0E-06 |
| KEGG:04141 | Processing in endoplasmic reticulum | 82 | 5.9E-05 |
| KEGG:03018 | RNA degradation | 40 | 4.6E-04 |
| KEGG:03050 | Proteasome | 35 | 5.5E-04 |
| KEGG:04144 | Endocytosis | 74 | 5.6E-04 |
| KEGG:04310 | Wnt signaling pathway | 24 | 1.8E-03 |
| KEGG:00071 | Fatty acid degradation | 24 | 2.9E-03 |
| KEGG:04130 | SNARE vesicular transport | 17 | 2.9E-03 |
| KEGG:00330 | Arginine and proline metabolism | 34 | 4.6E-03 |
| KEGG:00230 | Purine metabolism | 40 | 1.0E-02 |
| KEGG:03060 | Protein export | 17 | 1.1E-02 |
| KEGG:00280 | Valine, leucine and isoleucine degradation | 22 | 1.3E-02 |
| KEGG:00250 | Ala, asp and glu metabolism | 20 | 1.4E-02 |
| KEGG:00020 | Citrate cycle (TCA cycle) | 27 | 2.5E-02 |
| KEGG:00030 | Pentose phosphate pathway | 16 | 2.8E-02 |
| KEGG:04068 | FoxO signaling pathway | 16 | 2.9E-02 |
| KEGG:00500 | Starch and sucrose metabolism | 20 | 4.2E-02 |
| KEGG:03010 | Ribosome | 83 | 4.3E-02 |
| KEGG:03015 | mRNA surveillance pathway | 41 | 4.4E-02 |

\*Bonferroni Corrected

**Figure 2.**
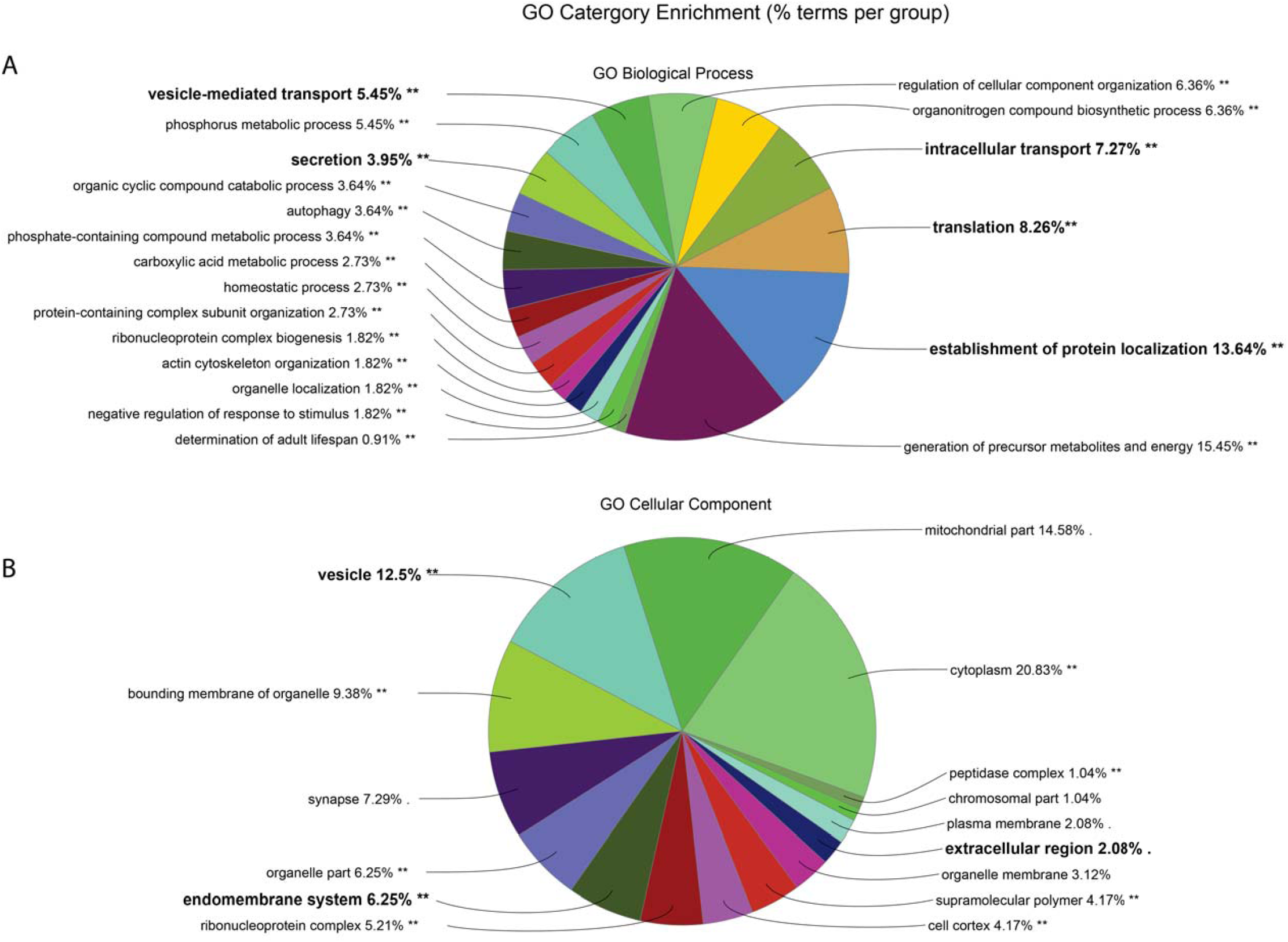
AcgP GO category enrichment analysis. Distribution of major functional category groupings for Biological Process (A) and Cellular Component (B). Categories related to secretory processes and discussed in main text highlighted in bold.

#### The Dpse accessory gland secretome (AcgS)

Signal peptides are a ubiquitous class of short (20-22 aa) N-terminal sequences, that target proteins for translocation across, and into, the endomembrane system of the cell (58, 59). Collectively proteins containing signal peptide sequences are considered part of the secretory pathway, and a subclass - those secreted into the extracellular space - are termed the secretome (also referred to as the exoproteome, exoP). Therefore, some or all components of the exoP can be considered as candidate SFPs. Given the secretory nature of the Drosophila accessory gland, we therefore queried the AcgP for proteins containing canonical signal sequences using two predictive programs, SignalP and Phobius (see Methods). SignalP (39) is a neural network-based algorithm designed to detect canonical N-terminal signal sequences and discriminate against N-terminal transmembrane regions known to reduce predictive power, and Phobius utilizes a combined model of both transmembrane and signal peptide predictors (40, 60). The combined output of both resulted in 771 Uniprot IDs that mapped to 535 Dpse FBgns (Figure 1B; Supplemental Table 3). Dmel orthologs (OrthoDB via Flybase) subsequently returned 506 Dmel orthologs to the Dpse AcgS including a small percentage (8/511, 1.6%) of “1:many” matches included in the analysis to capture the greatest proteome coverage of the secretome. Thus, the AcgS represents approximately 15% (535/3281; 16.3%) of the entire AcgP consistent with similar calculations for the predicted human secretome (∼15%, http://www.proteinatlas.org/humanproteome/secretome). Finally, the DRSC Integrative Ortholog Prediction Tools (DIOPT) website (http://www.flyrnai.org/diopt) identified 61% (327/535) of the AcgS that corresponded to orthologous human sequences (Supplemental Table 3).

### Gene Ontology (GO) Functional Analysis of the AcgS

The high degree of homology between the Dpse AcgS genes and Dmel (506/528), Supplemental Table 3) provided a putative orthologous secretome useful for GO analysis and network visualization. A significant enrichment in BP terms was observed, many related to multicellular organism reproduction (n=113, p=7E-7), reproduction (n=116, p=7E-7), behavior (n=53, p=2.1E-5 and proteolysis (n=66, p=3.5E-6). The secretome is enriched in MF terms related to oxidoreductase activity (n=55, p=7.1E-7) and hydrolase activity (n=138, p=8.5e-14; see Supplemental Table 3 for a complete list of all AcgS enriched BP, CC and MF GO categories). We also examined the subcellular localization of the AcgS using the Cerebral layout tool implemented in Cytoscape. As expected for functions related to secretion and proteins containing signal sequences targeted to the secretory pathway, the predicted subcellular localization of AcgS proteins were skewed toward extracellular and plasma membrane proteins (Figure 3).

**Figure 3.**
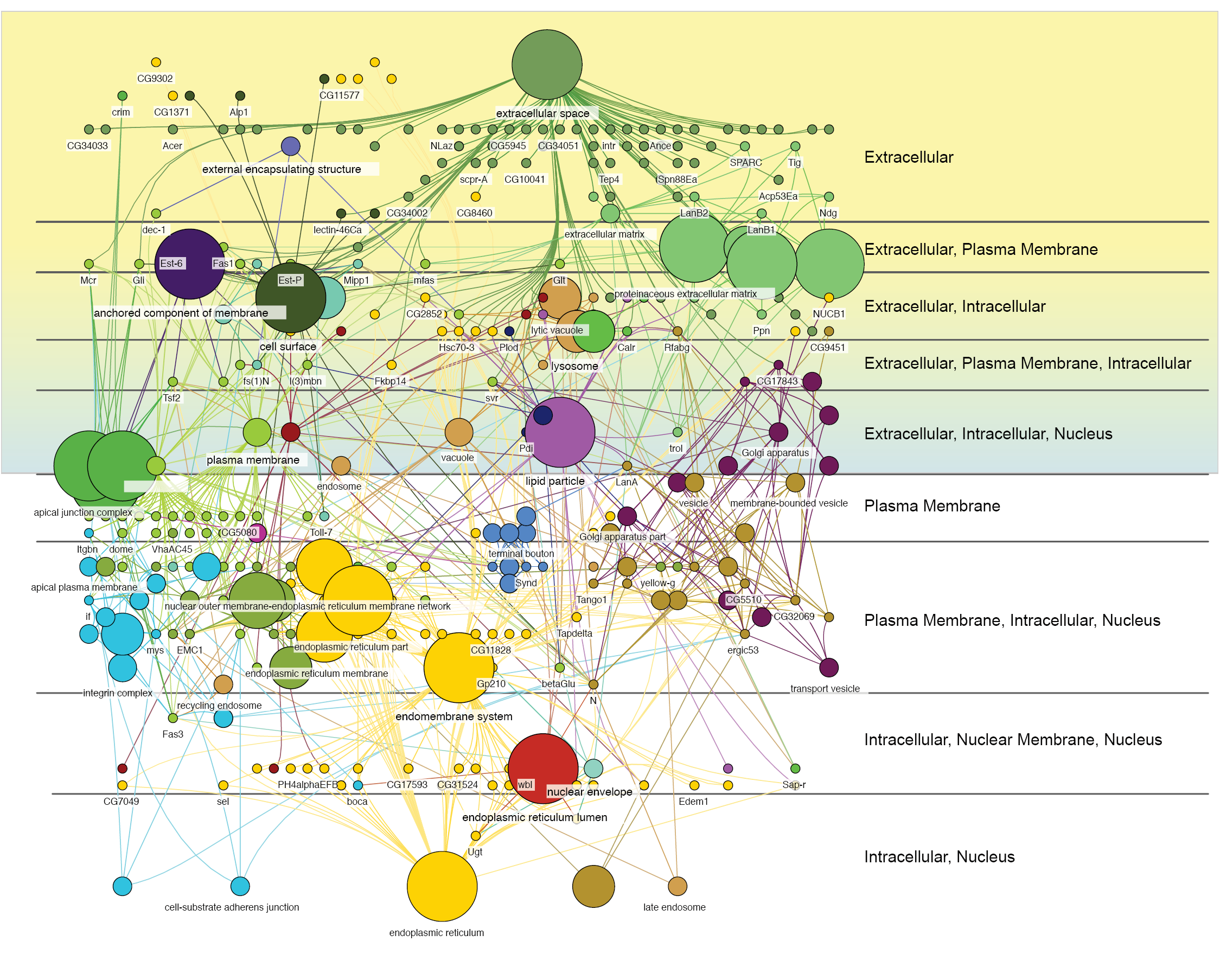
Functional grouping and subcellular localizaton of the AcgS mapped to subcellular regions (labeled on the right) using the Cerebral layout in Cytoscape. Shaded regions show all annotations that include the keyword “Extracellular”.

### The AcgS contains a robust repertoire of putative SFPs

We used the AcgS to identify a list of 151 putative SFPs representing >25% (151/506) of the AcgS (Figure 1B; Supplemental Table 4). This list is a conservative estimate as 99 genes had no CC functional annotation. Cytoscape and Cluego network analyses returned enriched BP terms of major functional categories including, insemination, sperm competition, and copulation reproduction, and negative regulation of endopeptidase activity (Table 2; Supplemental Table 4). Compared with a list of 212 Dmel SFPs assembled from the literature (9, 20, 21) an overlap of 32.1% (68/212) was observed that included 32 named SFP genes including Acp53Ea, Acp53C14d and Acp53C14c (Supplemental Table 4).

**Table 2.** Enriched BP categories of the AcgS exoproteome (putative SFPs).

| *BP Category* | *Nr. Genes* | *Pvalue\** |
| --- | --- | --- |
| multicellular organism reproduction | 77 | 7.5E-29 |
| insemination | 11 | 2.8E-13 |
| drug catabolic process | 10 | 1.3E-06 |
| response to biotic stimulus | 20 | 7.3E-06 |
| lipid transport | 9 | 2.2E-05 |
| response to wounding | 8 | 1.9E-03 |
| negative regulation of catalytic activity | 9 | 2.4E-03 |
| reproduction | 78 | 3.3E-27 |
| multi-multicellular organism process | 11 | 2.8E-13 |
| copulation | 11 | 2.7E-12 |
| sperm competition | 10 | 5.6E-12 |
| regulation of female receptivity, post-mating | 9 | 2.0E-11 |
| regulation of female receptivity | 9 | 1.7E-09 |
| female mating behavior | 9 | 2.0E-08 |
| response to external biotic stimulus | 20 | 7.3E-06 |
| response to other organism | 20 | 7.3E-06 |
| carbohydrate derivative catabolic process | 9 | 1.1E-05 |
| mating | 12 | 4.5E-05 |
| reproductive behavior | 12 | 6.2E-05 |
| mating behavior | 11 | 9.4E-05 |
| response to bacterium | 13 | 5.5E-04 |
| lipid localization | 9 | 9.6E-04 |
| aminoglycan metabolic process | 8 | 2.5E-03 |
| negative regulation of molecular function | 9 | 3.5E-03 |

\*Bonferonni corrected

### Divergent sexual selection results in differential AcgP protein abundance

To examine the proteome-wide response to sexual selection, differential protein abundance between the M-and P-lines was determined. A total of 250 differentially abundant Uniprot entries were identified using LFQ and Qspec statistical analysis across both treatments (FDR < 1%). The Uniprot IDs were subsequently mapped to 229 orthologous Dmel proteins (Supplemental Table 5). Taken together, network visualization of all 229 differentially abundant proteins revealed tight clustering of GO terms grouped into processes related to the cytoskeleton, protein translational machinery-particularly ribosomal proteins-and a suite of amino-acyl tRNA synthetases, along with a significant enrichment in categories related to endomembrane systems (e.g., Golgi, ER) and elements of the secretory pathway associated with vesicles and the Golgi apparatus (Figure 4A). Moreover, 14.4% (33/229) of the differentially abundant proteins were members of the exoP, including known Acps (Supplemental Table 6).

**Figure 4.**
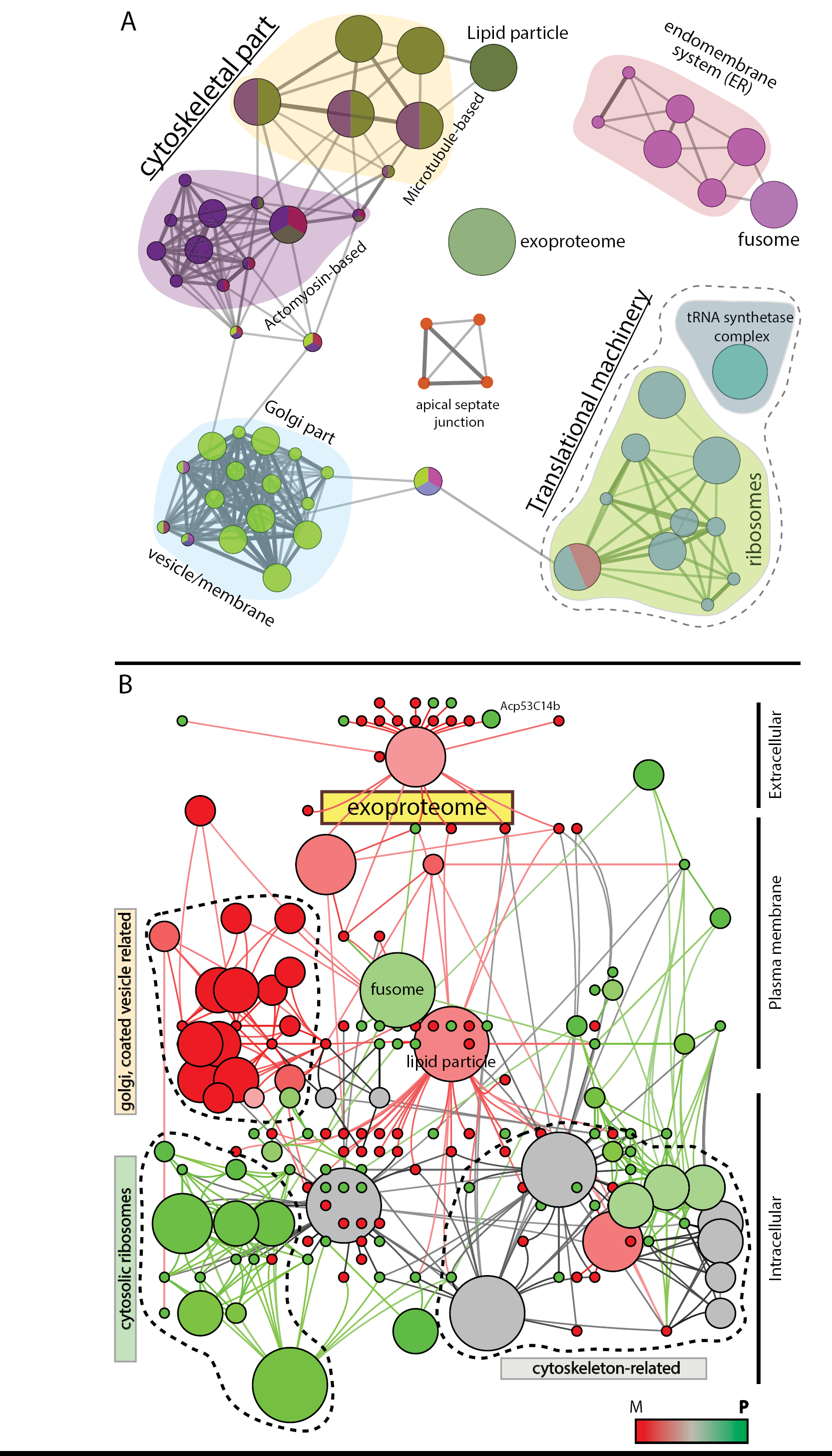
Network and predicted cellular distribution of DE proteins. (A) Overview of the DE proteins (M-and P-lines combined) showing major enriched functional groupings for GO Cellular Components (CC) using Cytoscape and Cluego. The four major groups, cytoskeletal part, golgi part, translation machinery and endomembrane system are shaded with different background colors. Dashed line encloses two functionally distinct categories, the ribosome and tRNA synthetase complex of the translation machinery (see also Table 5). (B) Network comparison of functional annotation groupings between DE proteins of the M-and P-lines. DE proteins were segregated into their respective M-or P-lines based on relative protein abundance and both groups compared using Cluego. A heat map (red-to-green) shows the distribution of DE protein between the M-and P-lines. These distributions were then mapped onto the predicted cellular locations of the groupings using the Cluepedia and the Cerebral layout as implemented in Cytoscape.

### M-and P-line differential protein abundance

To further probe the effect of PCSS on our divergent selection lines, we defined differential protein abundance between the sexual selection treatments based according to the sign of the log-fold change relative to the M-line (i.e., positive fold-change, M < P; negative fold-change, M > P; Supplemental Table 5). Of the 229 differentially abundant proteins, 133 and 96 were more abundant in the M-line in the P-line, respectively (Figure 1C). To examine differences in the function and cellular location of proteins differentially abundant between the M-and P-lines, the overlap of the two with respect to their GO groupings was generated using Cytoscape and Cluego. This comparison revealed a striking separation of functional grouping, with each selection line showing unique sub-groupings within the overall network of functional annotations (Figure 4B). We found differentially abundant proteins related to translational machinery (e.g., ribosomes) and cytoskeleton groupings predominant within the P-line (Figure 4B, green) with bias toward cytoplasmic cellular activities whereas Golgi-and vesicle-related groupings (including the exoP) biased in the M-line (Figure 4B, red) and functioning near or at the cell periphery.

Thirty-three of the differentially abundant proteins (24 up in M, 9 up in P) were annotated as “extracellular” (i.e., putative SFPs; Supplemental Table 6). While a larger number of candidate SFPs were more abundant in M-line compared to P-line males, over 50% of P abundant SFPs overlapped with known Dmel SFPs (5/9) compared to 29% overlap between M abundant and Dmel SFPs (7/24). Differentially abundant P male proteins did not show significant GO BP enrichment whereas significant enrichment was found in M-line males related to “response to fungus” (P = 2.1E-02) and “wound healing” (P = 2.0E-02) (9/24, 34.5%; Supplemental Table 6; Supplemental Figure 3).

### Sexual selection targets protein interaction networks

While the differentially abundant proteins of each selection line was biased towards distinct sets of GO functional groupings (Figure 4B), we also tested whether these relationships extended to specific protein interaction networks of known network topologies. We used the most stringent filters for network interactions on STRING (45), and found a statistically significant (P = 0.001) PPI network of 91 proteins containing nine defined subnetworks and including elements of 34 KEGG pathways (Supplemental Table 7). A striking aspect of the STRING-generated PPI networks was the almost complete segregation of M and P differentially abundant proteins into line-specific protein interaction networks (Figure 5). For example, all elements of the “vesicle transport” and “purine biosynthesis” PPI networks were more abundant exclusively in the M-line. Likewise, all proteins under “microtubule organization” and almost all (15/18) in the “ribosome” category are more abundant in the P-line. These results show that the divergent sexual selection treatments used to generate the M-and P-lines resulted in highly focused and interrelated changes in protein abundance that were specific to each treatment.

**Figure 5.**
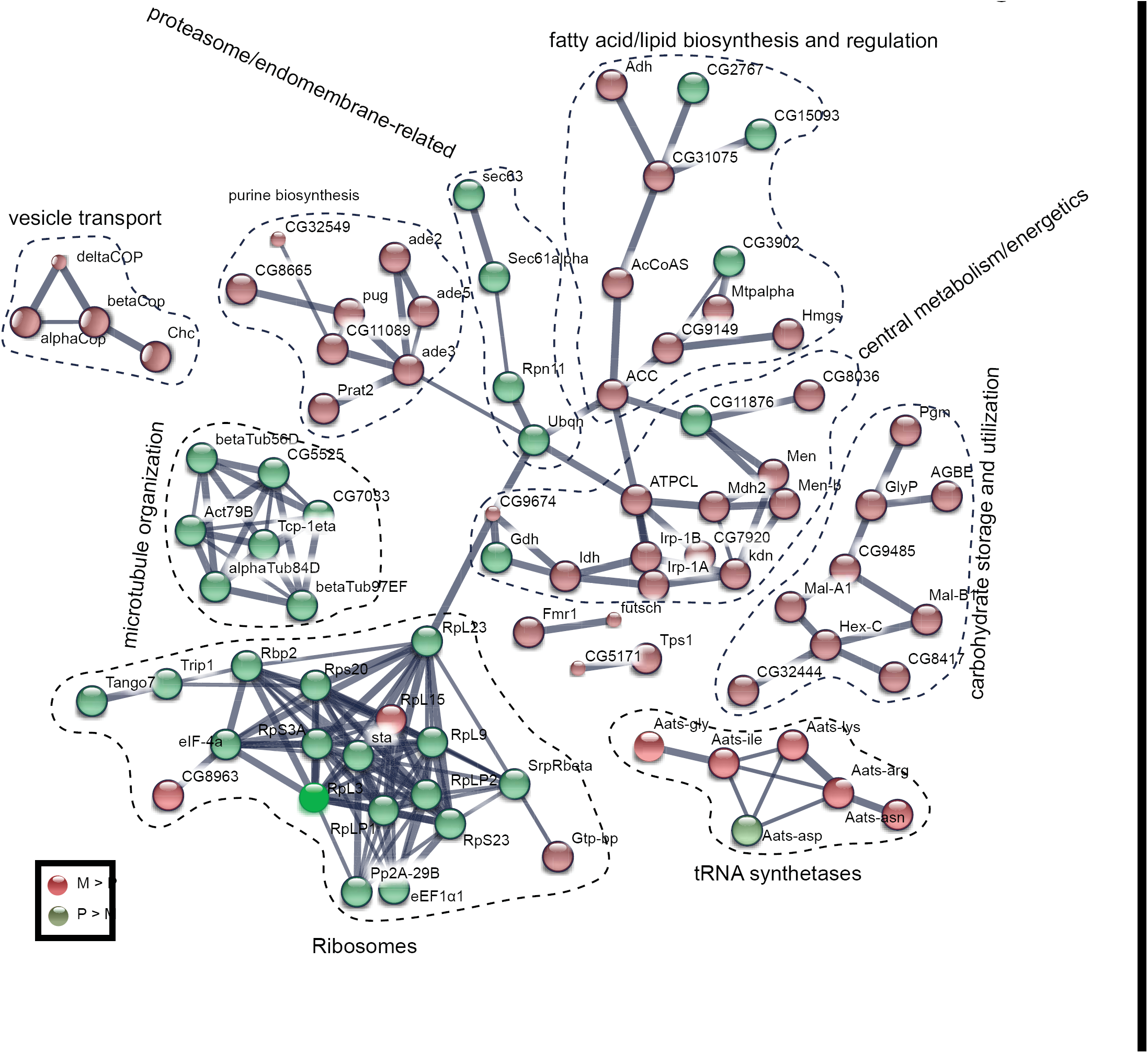
PPI network representation of the M (red, M>P) and P (green, P>M) DE proteins (inset). The PPI network was generated using STRING, a program that determines protein-protein interactions based on experimental and database criteria and calculated confidence levels to assign interactions between protein pairs. Stippled lines indicate major functional categories describing the networks.

### Molecular evolutionary rates of accessory gland protein genes

We tested for rates of molecular evolution in male reproductive proteins by estimating omega (dN/dS substitution rates) for each set of proteins: candidate SFPs, secretome proteins (minus SFPs), and the remaining accessory gland proteome proteins using orthologs from two closely related species in the obscura group, *D. lowei* and *D. affinis*. We found that putative Dpse SFPs (Exoproteome) are evolving faster than both accessory gland proteome proteins (AcgP; median omega = 0.088 versus 0.052, P=E-09) and accessory gland secretome minus the Exoproteome (AcgS; median omega = 0.088 versus 0.078, P = 2.7E-02; Figure 6A). AcgS proteins also evolve faster than the AcgP proteins (median omega = 0.052 versus 0.078, P= 1 × 10-9; Figure 6A).

**Figure 6.**
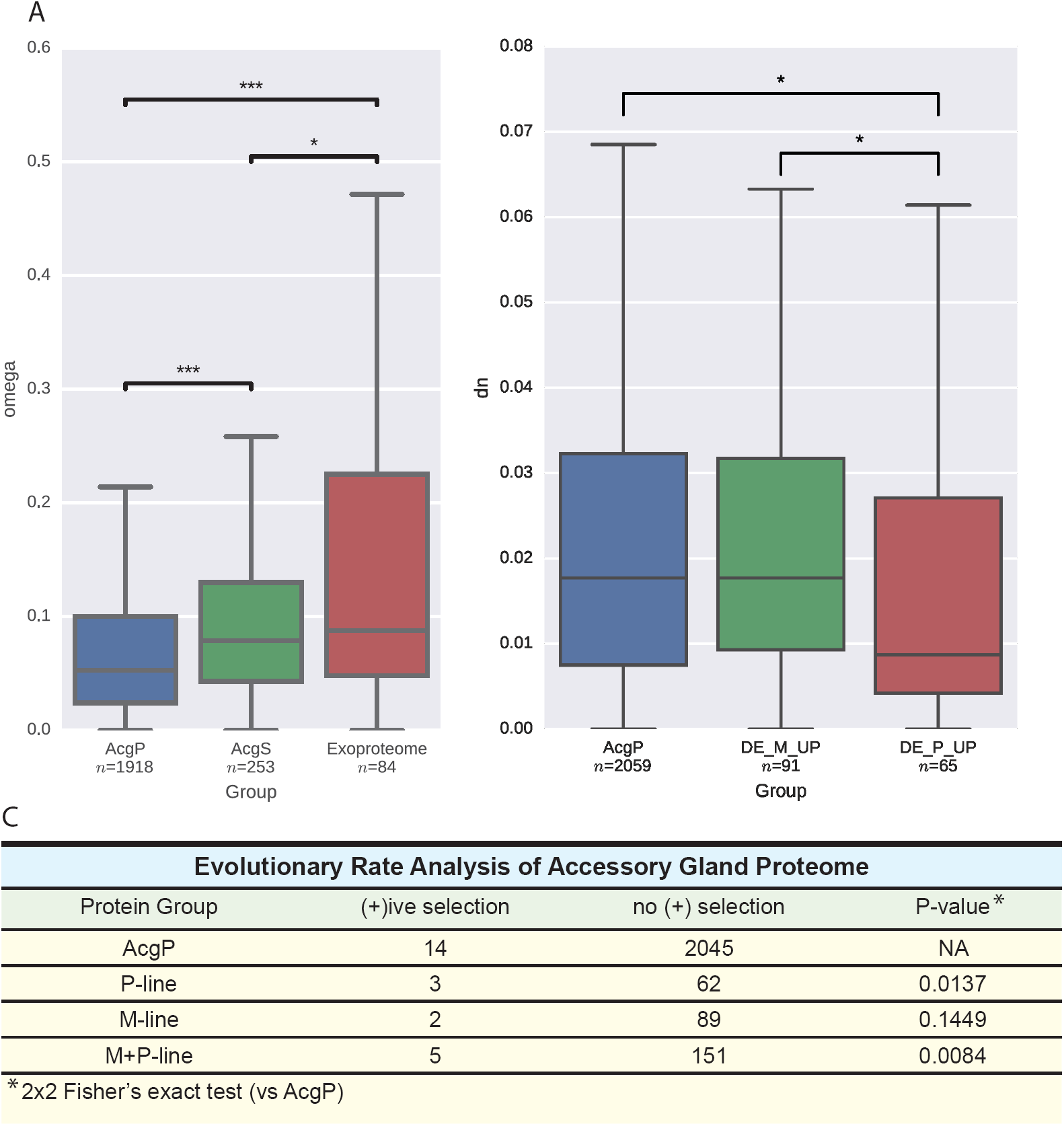
(A) Evolutionary rate calculations of the ratio of the non-synonymous to synonymous (dN/dS) substitution rates for (left to right) the Acg proteome (AgcP), the secretome (AcgS, excluding exoproteome and the exoproteome (AcgS-Extracellular). (B) The rate of nonsynonynous changes (dn) in genes coding for (left to right); the AcgP (minus DE M and P treatment genes), differentially abundant M-line genes (DE_M_UP) and differentially abundant P-line genes (DE_P_UP). Number of genes analyzed in each category listed below category. (C) Summary table of evolutionary rate analysis indicating significant enrichment of differential abundance proteins under positive selection.

Given that sexual selection is hypothesized to result in rapid protein sequence evolution, we then compared the evolutionary rates of the selection-specific differentially abundant proteins and accessory gland proteins (excluding selection-specific proteins). We found that the P treatment showed pronounced reduction in dN compared to the M treatment (Figure 6B, dN=0.009 (P) versus 0.018 (M), P=0.011). This is perhaps unsurprising considering that cytosolic ribosomal proteins are highly enriched in P, a protein group that has been shown to be slowly evolving (61). However, slowly evolving genes may be indicative of intensified selective pressures which obscure signals of positive selection, so to account for this we employed a direct site-specific test of positive selection across the accessory gland proteome. We identified 19 proteins that showed positive selection, 14 from the AcgP and 5 showing differential abundance in the selection lines, 3 in P and 2 in M (Table 3). An enrichment analysis shows that genes encoding proteins that were differentially abundant following experimental microevolution were more likely to be under positive selection than those in the AcgP set (Figure 6C, Supplemental Table 8) after correction for multiple testing and this is significant for the P-only set as well.

**Table 3.**
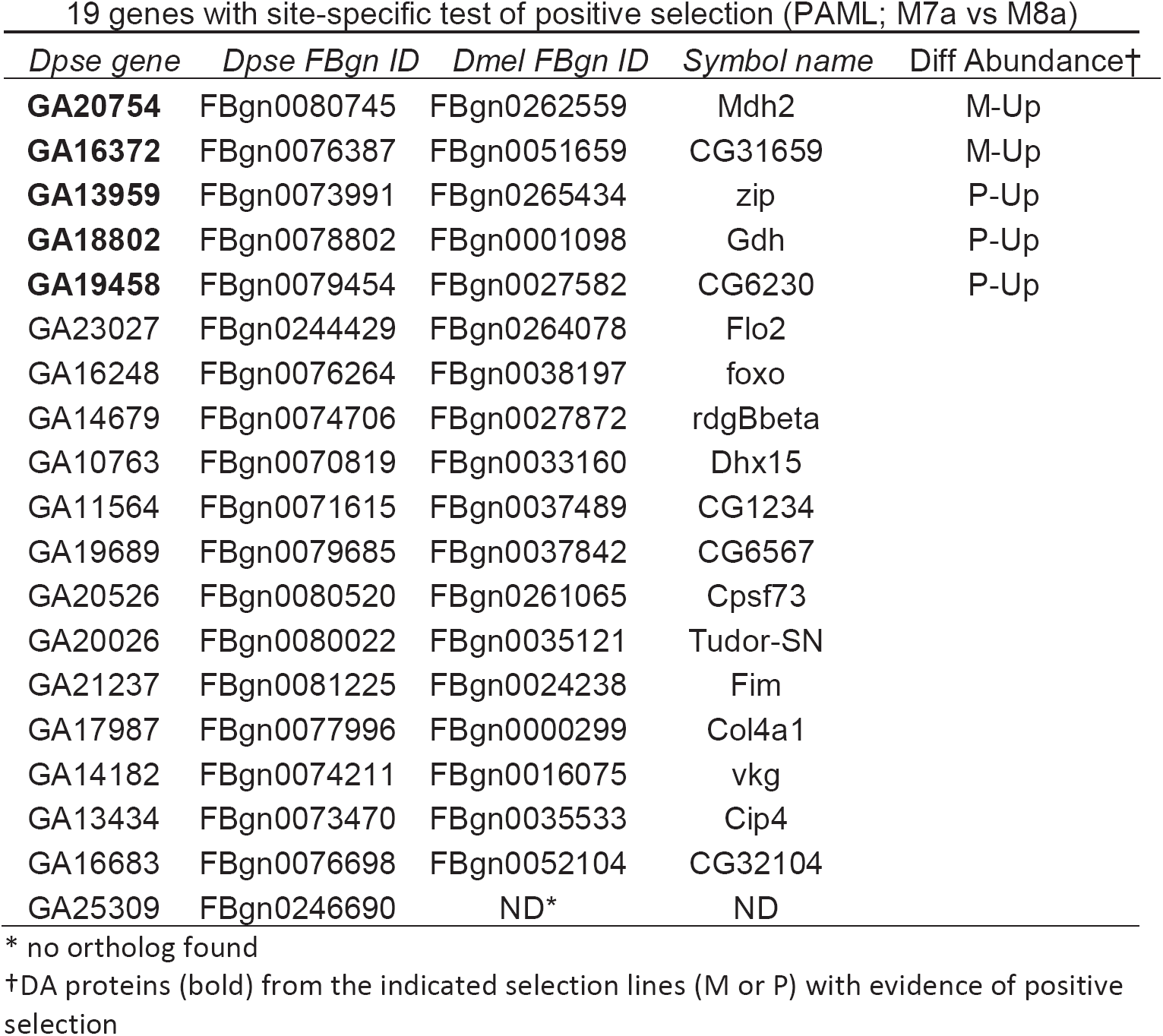
19 genes with site-specific test of positive selection (PAML; M7a vs M8a)

| *Dpse gene* | *Dpse FBgn ID* | *Dmel FBgn ID* | *Symbol name* | Diff Abundance† |
| --- | --- | --- | --- | --- |
| **GA20754** | FBgn0080745 | FBgn0262559 | Mdh2 | M-Up |
| **GA16372** | FBgn0076387 | FBgn0051659 | CG31659 | M-Up |
| **GA13959** | FBgn0073991 | FBgn0265434 | zip | P-Up |
| **GA18802** | FBgn0078802 | FBgn0001098 | Gdh | P-Up |
| **GA19458** | FBgn0079454 | FBgn0027582 | CG6230 | P-Up |
| GA23027 | FBgn0244429 | FBgn0264078 | Flo2 |  |
| GA16248 | FBgn0076264 | FBgn0038197 | foxo |  |
| GA14679 | FBgn0074706 | FBgn0027872 | rdgBbeta |  |
| GA10763 | FBgn0070819 | FBgn0033160 | Dhx15 |  |
| GA11564 | FBgn0071615 | FBgn0037489 | CG1234 |  |
| GA19689 | FBgn0079685 | FBgn0037842 | CG6567 |  |
| GA20526 | FBgn0080520 | FBgn0261065 | Cpsf73 |  |
| GA20026 | FBgn0080022 | FBgn0035121 | Tudor-SN |  |
| GA21237 | FBgn0081225 | FBgn0024238 | Fim |  |
| GA17987 | FBgn0077996 | FBgn0000299 | Col4a1 |  |
| GA14182 | FBgn0074211 | FBgn0016075 | vkg |  |
| GA13434 | FBgn0073470 | FBgn0035533 | Cip4 |  |
| GA16683 | FBgn0076698 | FBgn0052104 | CG32104 |  |
| GA25309 | FBgn0246690 | ND* | ND |  |

\* no ortholog found
†DA proteins (bold) from the indicated selection lines (M or P) with evidence of positive selection

## DISCUSSION

### Proteomics of the Dpse male accessory gland

The main aim of this work was to directly test the role of postcopulatory sexual selection in the evolution of reproductive proteins and the male accessory gland reproductive protein network. To accomplish this aim, we employed a “bottom up” shotgun proteomics approach, generating a robust accessory gland proteome containing 3281 proteins, representing the first accessory gland proteome to be described in Drosophila. The AcgP proteins in Dpse were enriched for cellular components expected from a tissue whose primary function is secretory, including several cellular component GO terms related to membranes, extracellular regions, and peptidase complex. The top biological process GO terms clearly indicated a large investment in processes directly and indirectly related to protein synthesis, protein assembly, transport and secretion. We then *in silico* concatenated this list to include proteins with secretory signal sequences to identify 545 accessory gland secretome proteins, which were enriched for GO terms related to the biological processes of reproduction, behavior and proteolysis, with these proteins heavily biased towards subcellular localizations in the plasma member or extracellular components, as predicted from proteins with secretory signals.

### Candidate Dpse SFPs contains many known Dmel SFPs

SFPs are secreted from the accessory gland tissue and transferred to the females upon mating. To generate a list of putative SFPs, we restricted the AcgS to only those proteins that were associated with GO cellular component annotations containing the keyword “extracellular”. The list of these proteins is most probably conservative as not all proteins were annotated. Nonetheless, the approach identified 151 proteins enriched for biological processes associated with reproduction, mating, insemination, and sperm competition, with approximately 30% that shared homology with previously identified Dmel SFPs. We also compared our list with the 29 putative SFPs previously identified via genome comparison between Dmel and Dpse (9). Of the nearly 50% overlap between these two lists (14/29), protein members of the canonical Sex Peptide (SP) network are particularly well-represented. In Dmel, SP binds to sperm in the female seminal receptacle, a sperm storage organ, and is required for both long-term female resistance to remating and for sperm release from storage (62). Our list of SFPs contains several known proteins involved in the Dmel SP network, including the gene duplicate pair lectins CG1652 and CG1656, aquarius (CG14061), intrepid (CG12558), antares (CG30488), seminase (CG10586), CG17575 (a cysteine-rich secretory protein), and CG9997 (a serine protease homolog) (16). CG9997 is processed in the female and males that do not produce this protein are unable to transfer the lectins, which are required to slow the rate at which CG9997 is processed in the female. All SP pathway proteins, except SP, were detected in our putative list of SFPs. Absence of detectable Dpse SP protein is consistent with the lack of a recognized SP ortholog in this species, and raises the interesting possibility that either the Dpse SP ortholog has significantly diverged, or has been replaced by another gene. If indeed a bona fide Dpse SP gene exists, further MS searches using algorithms to detect amino acid replacements (63) may be useful in the search for this elusive SFP.

### Functional compartmentalization results from divergent sexual selection

While our analysis found many known SFPs, 68% had no known orthology to Dmel This supports the idea that SFPs and related proteins involved in reproduction show rapid evolution as suggested by genome comparisons (9, 16, 17, 21, 64–66) and experimental evolution research on changes in sex-specific gene expression (32, 33, 67). Here we directly measured, using LC-MS/MS and LFQ, the proteome-wide effect of postcopulatory sexual selection on protein abundance in two populations that had undergone over 150 generations of divergent PCSS. This approach identified significant protein abundance differences in over 200 proteins between the selection lines. Knowledge of the changes in protein abundance under these experimental conditions provided a useful database to compare and contrast the effects of selection from a proteome-wide perspective and to assess the impact of selection on the protein-protein interaction networks revealed by our analysis.

Previous comparative work has suggested protein evolution may be adaptively compartmentalized for core processes and undergoing purifying selection and for interactive proteins that undergo positive selection (24, 26). We directly tested this assertion by comparing protein interaction network architecture in the divergent PCSS treatments and testing for molecular signatures of selection. We found remarkably line-specific compartmentalized changes to proteins after more than 150 generations of experimental sexual selection. P males invest in protein production machinery (i.e., protein quantity) and M males invest in protein surveillance (i.e., protein quality). The number of matings a Drosophila male can have is limited by SFP supply and how rapidly males can replenish this supply (68). Moreover, males can respond plastically to increased risk of sperm competition by transferring more of some SFPs to females (14, 69). We have previously shown that P males have larger accessory glands and can mate with more females sequentially than M males (30, 31). The increased investment in protein production machinery in P males may underlie this phenotypic response, and we suggest this may explain how males in polyandrous mating systems can rapidly adjust and replenish SFP quantity. In contrast, M males overall have increased production of proteins that function in downstream processing of secreted proteins, suggestive of increased investment in protein quality. The transfer of high quality proteins to females may be a response to reduce sexual conflict and improve population fitness, as predicted under enforced monogamy in which the reproductive interests of the sexes become aligned.

The SFPs produced in larger quantities in M compared to P males were enriched for chitin catabolic process (including wound healing) and response to fungus. Mating in Dmel initiates an immune response in females that has negative fitness consequences (70). The immune system may be activated to combat pathogens in the ejaculate and/or immunogenic sperm and SFPs (response to fungus), and/or wounds inflicted by males (chitin catabolic processes and wound healing). As enforced monogamy relaxes sexual conflict between males and females, M males may produce more of these proteins to limit damage to their mates. Future work could test this by manipulating these protein levels in the male ejaculate and determining the consequences on female fitness following mating. M males also produced more ejaculatory bulb protein (Ebp; CG2668), a known constituent of the mating plug. The mating plug remains in the female uterus until females eject it, along with unstored sperm. Ebp is necessary for ejaculate retention; knocking down Ebp in Dmel results in ejaculate loss, and reduced female sperm storage (Avila et al. 2015b). M males may produce more Ebp for better ejaculate retention, reducing the likelihood of their mates requiring additional matings to maintain sperm stores and subsequent fertility. Given the costs of mating for females, such a response is predicted when sexual conflict is relaxed, as in the M treatment.

Differentially abundant P male proteins did not show significant GO BP enrichment but over 50% of these (5/9) are known Dmel SFPs. This includes another mating plug protein, Acp53C14b, known to be involved in egg laying and reproductive fitness (71). Our previous work on the Dpse selection lines described sexual conflict over female reproductive phenology; M females mated to P males oviposit more eggs earlier in their lifespan than when mated to their own males (31). Given that the M treatment removes sexual conflict, alterations to the M female oviposition schedule when mated to P males is likely to be costly. This manipulation of female oviposition may be mediated by Acp53C14b given that this protein – which P males produce more of - influences oviposition.

### Rates of molecular evolution on male reproductive proteins

Rapid evolution at the molecular level is common for reproductive proteins including SFPs (e.g., (10, 18, 22, 26)) and this rapid evolution at the macroevolutionary scale is attributed to PCSS. Here we go beyond these traditional tests and ask whether there are differences in the rate of evolution between different types of reproductive proteins arising from the same tissue at the microevolutionary scale following over 150 generations of experimental manipulation of PCSS. We find that genes encoding SFPs showed significantly faster rates of molecular evolution compared to both other secreted protein encoding genes that were not candidate SFPs and overall accessory gland proteome genes. However, we also found that secretome genes showed higher rates of molecular evolution than non-secreted proteins (i.e., the remainder of the AcgP). These results support the interpretation that genes which interact extracellularly evolve more rapidly than those that remain within the cytoplasm (72, 73).

Evolutionary rates of genes encoding differentially abundant proteins between the sexual selection treatments found pronounced reduction in dN of P males compared to either differentially abundant proteins in M or the AcgP. This was likely due to the compartmentalization we identified, in which P males invested more in slowly evolving cytosolic ribosomal proteins (Barreto and Burton 2013). However, after controlling for this, we found that genes encoding for M-and P-differentially abundant proteins were more likely to have been subjected to positive selection compared to the AcgP genes as a whole. This result reinforces the hypothesis that the proteins we identified as responding to divergent sexual selection t the microevolutionary level have undergone adaptive processes on a macroevolutionary scale. We argue this directly illustrates a major role of PCSS in the evolution of male reproductive proteins.

## Conclusions

By employing divergent postcopulatory sexual selection through the use of experimental evolution of mating systems, we were able to directly test the hypothesis that postcopulatory sexual selection results in the rapid evolution of male-specific reproductive proteins. Within 150 generations, substantial microevolutionary changes occurred such as differential abundance of secreted proteins, including previously identified Dmel SFPs that have known effects on male and female reproductive fitness. A small number of these proteins showed signatures of positive molecular evolution. We found PCSS mediates remarkable compartmentalization of subcellular function of the secretory tissue, suggesting sexual selection alters fundamental reproductive protein networks, not just individual SFPs. These changes affected protein production, but in different ways in the divergent treatments, suggesting that PCSS selects for increased protein production whereas as relaxation of sexual conflict selects for protein surveillance. Some specific changes may be related to previously described phenotypic responses to divergent sexual selection that facilitate greater reproductive capacity in males subject to polyandry and ameliorates sexual conflict in populations subject to monandry. Our novel results suggest wide-spread, previously unappreciated, consequences of postcopulatory sexual selection on reproductive protein evolution. Combining the increasing use of high throughput proteomics and experimental manipulation of mating systems in different taxa will allow broader tests of this pattern in different taxa and better understanding of how sexual selection and sexual conflict impacts male reproductive protein evolution.

## Acknowledgements

We thank the many people who have contributed to the maintenance of the Snook experimental evolution lines. Funding of the experimental evolution lines came from NSF (DEB-0093149), NERC (NE/B504065/1; NE/I014632/1), and EU (ITN-2008-213780 SPECIATION) to RRS. Funding and technical support for the proteomic work came from the University of Sheffield Biological Mass Spectrophotometry Facility (funded by Yorkshire Cancer Research (Shend01) and the Wellcome Trust). TIG was supported by a Leverhulme Early Career Fellowship (Grant ECF-2015-453) and a NERC grant (NE/N013832/1). Funding from the Kyoto Institute of Technology to TLK also supported this research.

## DATA AVAILABILITY

Raw files for the entire datasets used in this study have been deposited via ProteomeXchange on the PRIDE database repository.

